# Screening and isolation of protease-producing bacteria from wastewater samples in Obafemi Awolowo University (OAU) Campus, Ile-Ife, Nigeria

**DOI:** 10.1101/2023.07.09.548249

**Authors:** Arogundade Femi Qudus, Lawal Ridwan Abiodun

**Author notes:** Corresponding Author: Arogundade Femi Qudus.

## Abstract

**Background:** Wastewater samples possess substantial potential as a valuable resource for the isolation of bacteria with the capacity to produce protease enzymes. Gaining insights into the proteolytic capabilities of these bacteria holds considerable significance for a wide range of industrial applications. Enhancing our understanding of the microbial diversity and protease production potential within wastewater can pave the way for the creation of customized enzymatic solutions tailored specifically for industrial needs.

**Purpose:** The purpose of this study was to isolate and identify protease-producing bacteria from wastewater samples collected at Obafemi Awolowo University Campus in Nigeria. The study involved isolating bacteria from the wastewater, identifying them, evaluating their growth on protease-supporting agar, determining their proteolytic activities, and screening bacterial colonies for protease production using skim milk agar medium.

**Methods:** Wastewater samples were aseptically collected from various locations within Obafemi Awolowo University Campus, located in Ile-Ife, Osun State, Nigeria. Bacterial isolation from the wastewater was performed using the serial dilution technique. The samples were progressively diluted and plated onto nutrient agar media for bacterial growth. Skim milk agar media was specifically used to isolate protease-producing bacteria. Following the isolation, screening was conducted to identify potential protease-producing bacteria. Zonal inhibition methods were employed using skim milk agar media during the screening process. The objective was to select bacterial isolates that exhibited clear zones around their colonies, indicating protease activity. To identify the potential protease-producing bacteria, morphological and biochemical tests were conducted. These tests included observations of colonial morphology, cellular morphology, and biochemical characteristics. The Bergey’s Manual was used as a reliable reference for taxonomic classification during the identification process.

**Results:** Among the ten bacterial colonies obtained from the wastewater samples, eight exhibited clear zones, indicating protease activity. Morphological and biochemical tests identified the protease-producing bacteria as *Bacillus spp*. and *Pseudomonas spp*. Further characterization revealed that the *Bacillus licheniformis* isolate from Water Sample D1 (WSD1) displayed the highest protease activity. *Bacillus subtilis* isolates also showed significant protease production, while *Pseudomonas spp*. exhibited lower protease production.

**Conclusions:** Wastewater samples from the OAU Campus yielded protease-producing bacteria, with *Bacillus licheniformis* showing the highest activity. The findings highlight the industrial potential of the isolated *Bacillus licheniformis* strain and emphasize the significance of utilizing wastewater as a source for obtaining bacteria with protease production capabilities. Further studies on individual strains within the *Bacillus* and *Pseudomonas* genera may lead to the discovery of strains with enhanced protease production, enabling tailored enzymatic solutions for various industrial sectors. Overall, this study contributes to our understanding of protease-producing microorganisms.

## 1.0 Introduction

Microbial enzymes play a crucial role in the food industry due to their superior stability compared to plant and animal enzymes. They can be efficiently produced through fermentation techniques, which offer cost-effectiveness, shorter production time, and reduced space requirements. The high consistency of microbial enzymes allows for easy process modification and optimization (Gurung et al., 2013).These enzymes have wide-ranging applications in various industrial sectors, as highlighted by Raveendran et al. (2018). One such enzyme, protease, finds utility in brewing, meat tenderization, coagulation of milk, and improvement of bread quality (Aruna et al., 2014; Miguel et al., 2013; Pandey et al., 2000). Similarly, α-amylase, another microbial enzyme, demonstrates its versatility in the food industry. It is employed in baking, brewing, starch liquefaction, bread quality improvement, production of rice cakes, and clarification of fruit juice (Kumar, 2015; Rodríguez et al., 2006; Aiyer, 2005; van der Maarel et al., 2002).

Proteases, enzymes responsible for the hydrolysis of peptide bonds in proteins and polypeptides, have significant industrial applications (Barrett & McDonald, 1986). The global demand for protease enzymes has been steadily growing, with a compound annual growth rate (CAGR) of 5.3% between 2014 and 2019 (Raveendran et al., 2018). They are extensively utilized in detergent, pharmaceutical, and food industries and account for 60% of the industrial enzyme market (Singh et al., 2016). Proteases, also known as proteolytic enzymes, play a crucial role in various biological processes and find widespread use in diverse industries (Raveendran et al., 2018). They are prominently employed in detergent formulations, leather processing, food processing, waste management, and pharmaceuticals. Specifically, proteases are used in brewing, meat tenderization, coagulation of milk, and bread quality improvement (Raveendran et al., 2018; Aruna et al., 2014; Miguel et al., 2013; Patel et al., 2013).The ability of microorganisms to produce proteases has garnered significant attention due to their potential for cost-effective and eco-friendly production.

Proteases can be classified based on their origin, catalytic activity and nature of the reactive group in the catalytic site. Microbes are an attractive source of proteases owing to the limited space required for their cultivation and their ready susceptibility to genetic manipulation (Rao et al., 1998). The major sources of protease enzymes are animals, plant and microorganisms (both bacterial and fungal). Proteases are divided into two groups: exopeptidases and endopeptidases, based on the site of action on polypeptide chains (Rao et al., 1998). The endopeptidases are further classified into six groups, based on the catalytic residue present in the active site: serine, aspartic, cysteine, metallo, glutamic acid and threonine protease. The exopeptidases act on the ends of polypeptide chains and endopeptidases act randomly in the inner regions of polypeptide chains (Li et al., 2013).

Proteases can also be classified into three groups based on their acid-base behavior: acid, neutral, and alkaline proteases. Acid proteases are produced primarily by fungi and exhibit optimal activity at a pH range of 2.0-5.0. Neutral proteases, on the other hand, are mainly of plant origin and have an optimal pH range around 7.0. Alkaline proteases, which exhibit maximum activity at pH values of 8 and above, are produced by several microorganisms, including bacteria (Alnahdi, 2012). Alkalophilic proteases, in particular, have demonstrated significant utility in the detergent industry, while acidophilic proteases have found applications in leather tanning, food processing, and x-ray film production (Habib et al., 2012; Rayda et al., 2012; Badgujar & Mahajan, 2013). The genus *Bacillus* is vital for commercially important alkaline protease (EC.3.4.21-24.99), which is active at alkaline pH ranging between 9 and 11 (Varela et al., 1997; Kocher & Mishra, 2009; Singhal et al., 2012). These alkaline protease producers are distributed in water, soil, and highly alkaline conditions. Alkaline proteases are unique in their activity and maintain a constant alkaline pH while being exploited for different formulations in pharmaceutical, food, and other related industries (Banerjee et al., 1999; Joo et al., 2002, 2004; Dias et al., 2008). A broad range of applications of these alkaline proteases are getting more attention from researchers with the hope of discovering new strains with unique properties and substantial activity (Najafi et al., 2005; Saeki et al., 2007). Two essential types of alkaline proteases, such as subtilisin Carlsberg and subtilisin novo are obtained from *Bacillus spp*., which can be used as an industrial enzyme to produce zein hydrolysates (Miyaji et al., 2006). Acid proteases are stable and active between pH 3.8 and 5.6 and are frequently used in soy sauce, protein hydrolysate, and digestive aids and in the production of seasoning material. The optimum pH of acidic proteases is 3–4 and the isoelectric point range is between 3 and 4.5 with a molecular weight of 30–45 kDa (Zheng et al., 2011; Ravikumar et al., 2012; Machado et al., 2016). In comparison with alkaline proteases, these extracellular acid proteases are mostly produced by fungal species, such as *Aspergillus niger* (Sielecki et al., 1991), *<u>Aspergillus oryzae</u>* (Yongquan, 2001), *Aspergillus awamori* (Ottesen & Rickert, 1970), *Aspergillus fumigatus* (Shinmyo et al., 1972), and *Aspergillus saitoi* (Sodek & Hofmann, 1970). Acid proteases are also exploited for use in clearing beer and fruit juice, improving texture of flour paste, and tenderizing the fibril muscle (Zhang et al., 2010). Neutral proteases are defined as, such as they are active at a neutral or weakly acidic or weakly alkaline pH. Mostly neutral proteases belong to the genus *Bacillus* and with a relatively low thermotolerance ranging from pH 5 to 8. They generate less bitterness in hydrolysis of food proteins due to a medium rate of reaction; therefore, they are considered more valuable in the food industry. Neutrase is incorporated in the brewing industry due to its insensitivity to plant proteinase inhibitors (Sodek & Hofmann, 1970).

Since the advent of enzymology, microbial proteolytic proteases have been the most widely studied enzyme (Razzaq et al., 2019). These enzymes have gained interest not only due to their vital role in metabolic activities but also due to their immense utilization in industries (Rao et al., 1998; Sandhya et al., 2005; Younes & Rinaudo, 2015). The proteases available in the market are of microbial origin because of their high yield, less time consumption, less space requirement, lofty genetic manipulation, and cost-effectiveness, which have made them suitable for biotechnological application in the market (Nisha & Divakaran, 2014; Ali et al., 2016). Comparatively, proteases produced by plants and animals are more labor-intensive than microbially produced proteases (Gupta et al., 2002; Kalaiarasi & Sunitha, 2009). Microbial proteases are preferred to plant and animal proteases because of the presence of all desired characteristics for industrial applications (Palsaniya et al., 2012; Sathishkumar et al., 2015).

Protease producing microorganisms have been isolated from a variety of environments like rhizosphere soil, slaughter house soil or floor washing, sewage, food waste, etc. (Jadhav et al. 2020; Hakim et al. 2018; Ash et al. 2018; Prajapati et al. 2017). These have also been from saline environments like the marine environments, sea sediments, hypersaline lakes, salted food, soda lakes, etc. (Maruthiah et al. 2016; Boughachiche et al. 2016; Ibrahim et al. 2019). Wastewater represents a rich source of diverse microorganisms, including protease-producing bacteria. These bacteria have adapted to thrive in the complex and nutrient-rich environment of wastewater, where they encounter various organic compounds, including proteins. Consequently, the screening and isolation of protease-producing bacteria from wastewater samples offer an opportunity to discover novel microbial strains with robust proteolytic capabilities.

This study focused on screening and isolating protease-producing bacteria from wastewater samples collected from different locations within Obafemi Awolowo University (OAU) Campus in Ile-Ife, Nigeria. By utilizing wastewater as a sample source, the study aimed to identify bacterial strains that were well-adapted to the local environment and possessed desirable proteolytic properties. The core objectives of this study encompassed the isolation of protease-producing bacteria from wastewater, identification of the isolated microorganisms, assessment of their potential to grow on protease-supporting agar, determination of the proteolytic activities of the isolated microbes, and screening of bacterial colonies for protease production on skim milk agar medium.

## 2.0 Materials and Methods

This study was conducted at the Department of Microbiology, Obafemi Awolowo University, Ile-Ife, Osun State, Nigeria.

### 2.1 Equipment

A variety of equipment and glassware were utilized in the study, including a sensitive weighing balance, autoclave, hotplate, incubator, inoculating loop, refrigerator, foil paper, permanent markers, Petri dishes, spatula, Bunsen burner, cotton wool, conical flask, measuring cylinder, McCartney bottles, pipette, and test tubes. The media employed in the experiment were nutrient agar and skimmed milk agar, and the sample analyzed was wastewater.

### 2.2 Sample Collection

Aseptic collection of wastewater samples was carried out from various locations across Obafemi Awolowo University Campus, situated in Ile-Ife, Osun State, Nigeria. The samples were transported under refrigerated conditions and promptly analyzed at the microbiology laboratory within 24 hours of collection. All experimental procedures were conducted within the laboratory to ensure accuracy and precision of the results.

### 2.3 Preparation of Materials

#### 2.3.1 Cleaning and Sanitizing of the Work Bench

To ensure a sterile laboratory environment, the workbench was subjected to sanitation procedures employing 70% ethanol. Specifically, the surface was meticulously swabbed using cotton wool immersed in alcohol, facilitating the removal of any bacteria and non-bacterial particulate matter. This rigorous approach to sanitization minimized the potential for contamination and promoted optimal experimental conditions.

#### 2.3.2 Sterilization of Materials

To ensure the sterility of the laboratory equipment and materials, a comprehensive sterilization process was implemented. Measuring cylinders, pipettes, and beakers were subjected to a sterilization cycle in a hot air oven set at 180°C for a duration of 2 hours. This high-temperature treatment effectively eliminated any potential contaminants.

Culture media and reagents underwent sterilization using an autoclave operating at 120°C under a pressure of 15 psi for 15 minutes. The autoclave’s combination of high temperature and pressure created an environment that effectively sterilized the media and reagents, rendering them free from any microorganisms.

Specific materials like inoculating loops, needles, and the mouths of flasks containing culture media and test tubes were subjected to flame sterilization. The intense heat from the flame effectively eliminated any residual contaminants, ensuring the sterility of these critical components.

By implementing this rigorous sterilization protocol, the laboratory equipment and materials were rendered free from contaminants that could potentially compromise the accuracy and reliability of the experimental results.

#### 2.3.3 Media Preparation

In accordance with microbiological standard protocols, various growth media were prepared and their corresponding weights were accurately measured using a precision balance and transferred into conical flasks. Subsequently, the media were fully dissolved in distilled water and sterilized by subjecting them to 15 psi at 121°C for 15 minutes, as per the recommendations of Harrigan & McCance (1996).

##### 2.3.3.1 Nutrient Agar Preparation

As per the standardized method recommended by Harrigan & McCance (1996), 28 grams of nutrient agar powder were meticulously weighed and combined with 1000 milliliters of distilled water in a conical flask. The mixture was then stirred with a glass stirrer to ensure homogeneity, and subsequently subjected to sterilization through autoclaving at 121°C for 15 minutes. This rigorous sterilization process aimed to eliminate all forms of life that may be present in the media (Harrigan & McCance, 1996).

##### 2.3.3.2 Saline Skim Milk Agar Preparation

In accordance with standard methodology (APHA, 1998), a 100 mL conical flask was charged with a mixture of 2.5% (w/v) skimmed milk, 0.5% (w/v) peptone, 1.0% (w/v) NaCl, and 1.5% (w/v) agar, which was subsequently dissolved in distilled water. To ensure uniformity of the mixture, stirring was carried out with a glass stirrer while heating. The resulting mixture was sterilized by autoclaving at 121°C and 15 psi for 15 minutes, as recommended by APHA (1998).

### 2.4 Method of Isolation

The serial dilution technique was employed to decrease the initial microbial load present in the water sample. To isolate microorganisms from water samples, aseptic technique was employed. 1 mL of water sample was aseptically transferred into 9 mL of sterile distilled water and mixed properly. From this mixture, 1 mL was transferred into a second test tube labeled 10^-2^ containing 9 mL of sterile distilled water and mixed properly. This process was repeated for the remaining three test tubes containing 9 mL of sterile distilled water to obtain dilutions of 10^-3^, 10^-4^, and 10^-5^.

The Nutrient agar was prepared and allowed to cool to around 45°C. Using a sterile pipette, 1 mL of each 10^-^3 and 10^-^5 dilution was aseptically transferred onto separate agar plates, which were properly labeled. Approximately 15 mL of molten agar was poured onto each plate, ensuring even distribution, and gently swirled to facilitate the appropriate spacing of colonies. After pouring the agar, the plates were left undisturbed to solidify, and then incubated at a temperature of 37°C for a duration of 24 hours and observed for microbial growth.

#### 2.4.1 Isolation of Pure Organisms

The process of subculturing involved using a sterile inoculating loop to gently pick the colony of interest, ensuring that two different colonies were not picked together, and streaking them on a newly prepared media. In this study, nutrient agar was utilized for bacterial growth, and it was inoculated for 18-24 hours, following the procedure outlined by Olutiola et al. (1991).

### 2.5 Screening for the Protease Degrading Isolates

The experimental protocol involved the use of a pour plate technique (Sanders, 2012), which is a standard method in microbiology, to culture and isolate proteolytic microorganisms. A medium containing 2.5% (w/v) skimmed milk, 0.5% (w/v) peptone, 1.0% (w/v) NaCl, and 1.5% (w/v) agar was utilized in the study. The medium was prepared and sterilized appropriately before being poured onto sterile plates. The plates were subsequently inoculated with the test organisms and incubated at room temperature for 24-36 hours.

The medium was carefully formulated to provide essential nutrients for the growth of proteolytic microorganisms. Skimmed milk and peptone provided sources of protein, while NaCl served as an osmotic stabilizer, and agar was added as a solidifying agent. The use of this medium enabled the growth of microorganisms that are capable of utilizing protease substrate, which was detected by screening the growth of the isolates on the medium.

The pour plate technique facilitated the isolation of proteolytic microorganisms by enabling their growth on the surface of the medium. The ability of the organisms to grow on the medium was indicative of their proteolytic activity. Therefore, only the organisms that demonstrated the ability to utilize the protease substrate were able to grow on the medium.

### 2.6 Preservation of Isolates on Slants

The pure cultures of the isolates, preferably (18-24 hours old) that was subcultured were aseptically transferred to the already prepared nutrient agar slants. The slants were stored in the refrigerator and the preserved cultures were used for all the tests that were carried out on the isolates (Olutiola et al., 1991).

### 2.7 Characterization and Identification of Isolates

Following the screening for efficient proteolytic activity, colonies exhibiting promising results were subjected to bacterial identification studies, including morphological and biochemical tests. The bacterial isolates obtained from the samples were characterized based on their colonial morphology, which involved observing characteristics such as shape, colour, size, elevation, Gram reaction, and pigmentation. Cellular morphology was also examined, focusing on the microscopic appearance of the cells. In the identification process, the Bergey’s Manual, a recognized reference for bacterial taxonomy and classification, was utilized as a resource (Olutiola et al., 1991).

#### 2.7.1 Microscopic Examination of Isolates

The bacteria isolates were characterized based on the Gram stain techniques which apart from differentiating an isolate as Gram positive or negative, also helped to identify whether it is rod or cocci (Olutiola et al., 1991).

##### 2.7.1.1 Gram-Staining

After preparing a 24-hour-old culture, it was smeared on the slide and fixed with heat. Subsequently, it was treated with crystal violet for a duration of 2 minutes, followed by rinsing with running water. The slide was then flooded with gram iodine solution for 2 minutes, and washed with alcohol. Finally, it was flooded with safranin, washed with water, dried, and observed under oil immersion (Harley & Prescott, 2002).

#### 2.7.2 Biochemical Characterization of Bacterial Isolates

The bacteria were subjected to biochemical characterization by conducting specific tests, including catalase, oxidase, methyl red, indole, Voges-Proskauer, and citrate tests. These tests were performed to evaluate the bacteria’s metabolic capabilities and enzymatic activities.

##### 2.7.2.1 Catalase Test

This test detected the presence or absence of catalase in the isolates. Catalase is an enzyme found in most bacteria and is known to catalyze the breakdown of hydrogen peroxide to liberate molecular oxygen (O_2_).

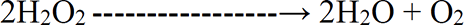

To a loopful of the culture, a few drops of 3% hydrogen peroxide were added on a slide, and any reaction was observed. The evolution of gas or white froth indicated a catalase positive reaction, while the absence of froth indicated a negative reaction (Boominadhan et al., 2009; Olutiola et al., 1991).

##### 2.7.2.2 Methyl Red Test

Ten milliliters of glucose phosphate peptone broth were dispensed into test tubes, which were then capped with cotton wool wrapped in foil and sterilized. The test organisms were inoculated and incubated at 37°C for 48 hours. After the incubation period, a few drops of methyl red indicator were added to the cultures. The production or development of a red colouration was considered a positive reaction, while the absence of colour indicated a negative reaction (Olutiola et al., 1991)

##### 2.7.2.3 Voges Proskauer Test

Sterile glucose phosphate broth (10ml) in a test tube was inoculated with the test isolates and incubated for 48 hours at 37°C. After incubation, 0.5ml of 0.5% alpha-Naphthol solution and 0.5ml of 16% potassium hydroxide solution were introduced into the mixture. The mixture was then shaken vigorously and left for about 10 minutes. A red colouration indicated a positive result, demonstrating acetoin production, while the absence of colouration indicated a negative result. This test demonstrated the production of acetyl methyl carbinol from glucose (Cheesebrough, 2002).

##### 2.7.2.4 Citrate Utilization

The preparation and sterilization of Simmons citrate agar tubes were completed, and all the isolates were then inoculated through stab inoculation. The tubes were subsequently incubated at 37°C for 24 hours. A positive result was observed as the indicator changed from green to an intense Prussian blue colour (Pandian et al., 2012). The colour change from green to Prussian blue indicates the utilization of citrate. This test was designed to assess the ability of microorganisms to utilize citrate as their sole source of carbon (Olutiola et al., 1991).

##### 2.7.2.5 Indole Test

The peptone broth medium was prepared and autoclaved. After cooling, the medium was inoculated and incubated at 30 °C for 48 hours, with a control kept for comparison. Kovac’s reagent was added gradually, drop by drop. The tubes were gently shaken and left to stand for 10 minutes to allow the formation of a distinct layer. The presence of a red-coloured layer on top of the tube indicated a positive result (Kiran et al., 2002; Olutiola et al., 1991).

##### 2.7.2.6 Sugar Fermentation Test

This was a test conducted to confirm the ability of bacterial isolates to utilize various sugars. A 1% (w/v) peptone water and sugar broth was prepared by combining peptone water and the respective sugars. Bromocresol was added as an indicator. The sugars included glucose, galactose, mannitol, fructose, lactose, etc. Each test tube was dispensed with 10 ml of the broth, with Durham tubes inverted inside them, and sterilized by autoclaving at 110°C for ten minutes. The test organisms were then inoculated into the broths in the test tubes and incubated at 37°C for 48 hours. Uninoculated tubes served as controls. A positive reaction was indicated by a colour change of the broth medium from red to yellow, and sometimes gas production in the Durham tube (Olutiola et al., 1991)

### 2.8 Growth and Physiological Characteristics

#### 2.8.1 Growth at Temperature 20^0^C and 70^0^C

Nutrient agar plates were prepared, and 18-24 hours old cultures of the bacterial isolates were inoculated onto the different plates. The plates were then incubated at high (70°C) and low (20°C) temperatures. The purpose of the test was to determine the temperature that favored the growth and metabolism of the bacteria, as indicated by their growth on the agar. Uninoculated plates were used as controls during the test.

#### 2.8.2 Growth In 6.5% Sodium Chloride (NaCl) Solution

An inoculum from a pure culture was aseptically transferred into a sterile tube containing 6.5% sodium chloride broth. The incubation was conducted at 35-37°C for 24 hours. If the microbes could grow in the broth, it became turbid after incubation.

#### 2.8.3 Growth in Different pH (pH 3.9 and pH 9.4)

Hydrochloric acid was added dropwise to homogenized molten Nutrient agar until a pH of 3.9 was reached for the acidic medium. Similarly, sodium hydroxide was added dropwise and homogenized for 15 minutes at 121°C before being poured into plates after cooling to approximately 45°C. The 24-hour-old bacterial isolates were streaked on the solidified agar and incubated at 37°C for 24 hours. This test helped determine the optimal pH for the growth of the isolates. Uninoculated plates served as control experiments.

## 3.0 Results

### 3.1 Mean Microbial Load of the Isolates

The mean microbial count of the waste water sample is presented in this Table 1. The total bacteria count (TBC) ranged from 1.9×10^5^ to 3.84×10^5^cfu/ml. While the total coliform count ranged from 3.53×10^2^ to 4.75×10^2^cfu/ml.

**Table 1:** Mean Microbial Load of Bacteria Isolates

| S/N | Sampling | MBC cfu/ml | TCC cfu/ml |
| --- | --- | --- | --- |
| 1 | 1st sampling | $2.4 \times 10^5$ | $3.23 \times 10^5$ |
| 2 | 2nd sampling | $1.9 \times 10^5$ | $4.58 \times 10^5$ |
| 3 | 3rd sampling | $3.5 \times 10^5$ | $3.42 \times 10^5$ |
| 4 | 4th sampling | $4.5 \times 10^5$ | $4.98 \times 10^5$ |
| 5 | 5th sampling | $5.5 \times 10^5$ | $3.66 \times 10^5$ |

**KEY:** MBC-Mean Bacteria Count TCC- Total Coliform count

### 3.2 Colony Morphology of the Bacteria Isolates

The morphological characteristics of bacteria isolates are presented in Table 2. The bacteria isolated showed different cultural characteristics, some colony appeared yellow and creamy in colour, some were circular and irregular in shape, all had surfaces which were smooth, most had margins that were entire and others undulate and lobate, different elevations such as smooth and flat and none of the organisms produced pigment.

**Table 2:** Colony Morphology of Bacteria Isolated from the Samples

| S/N | Isolates | Shape | Colour | Elevation | Edge | Size | Surface | Margin |
| --- | --- | --- | --- | --- | --- | --- | --- | --- |
| 1 | WSA1 | Irregular | Yellow | Raised | Entire | Large | Smooth | Entire |
| 2 | WSA5 | Circular | Cream | Raised | Undulate | Small | Smooth | Undulate |
| 3 | WSA6 | Irregular | Orange | Flat | Entire | Small | Rough | Entire |
| 4 | WSA7 | Irregular | Cream | Raised | Entire | Small | Smooth | Entire |
| 5 | WSA8 | Circular | Cream | Flat | Entire | Small | Wet | Entire |
| 6 | WSD1 | Circular | Yellow | Raised | Entire | Small | Smooth | Entire |
| 7 | WSD4 | Circular | Yellow | Raised | Entire | Large | Smooth | Entire |
| 8 | WSE4 | Circular | Cream | Flat | Entire | Small | Rough | Entire |
| 9 | WSF1 | Circular | Orange | Raised | Entire | Small | Smooth | Entire |
| 10 | WSF5 | Irregular | White | Flat | Entire | Small | Rough | Entire |

### 3.3 Observation of Clear Zone on Skim Milk Agar

The bacteria isolated showed zone of inhibition on skim milk agar at 37^0^C for 48 hours. The zone of hydrolysis was noted for each sample. Table 3 shows zone of inhibition of isolated bacteria.

**Figure 1:**
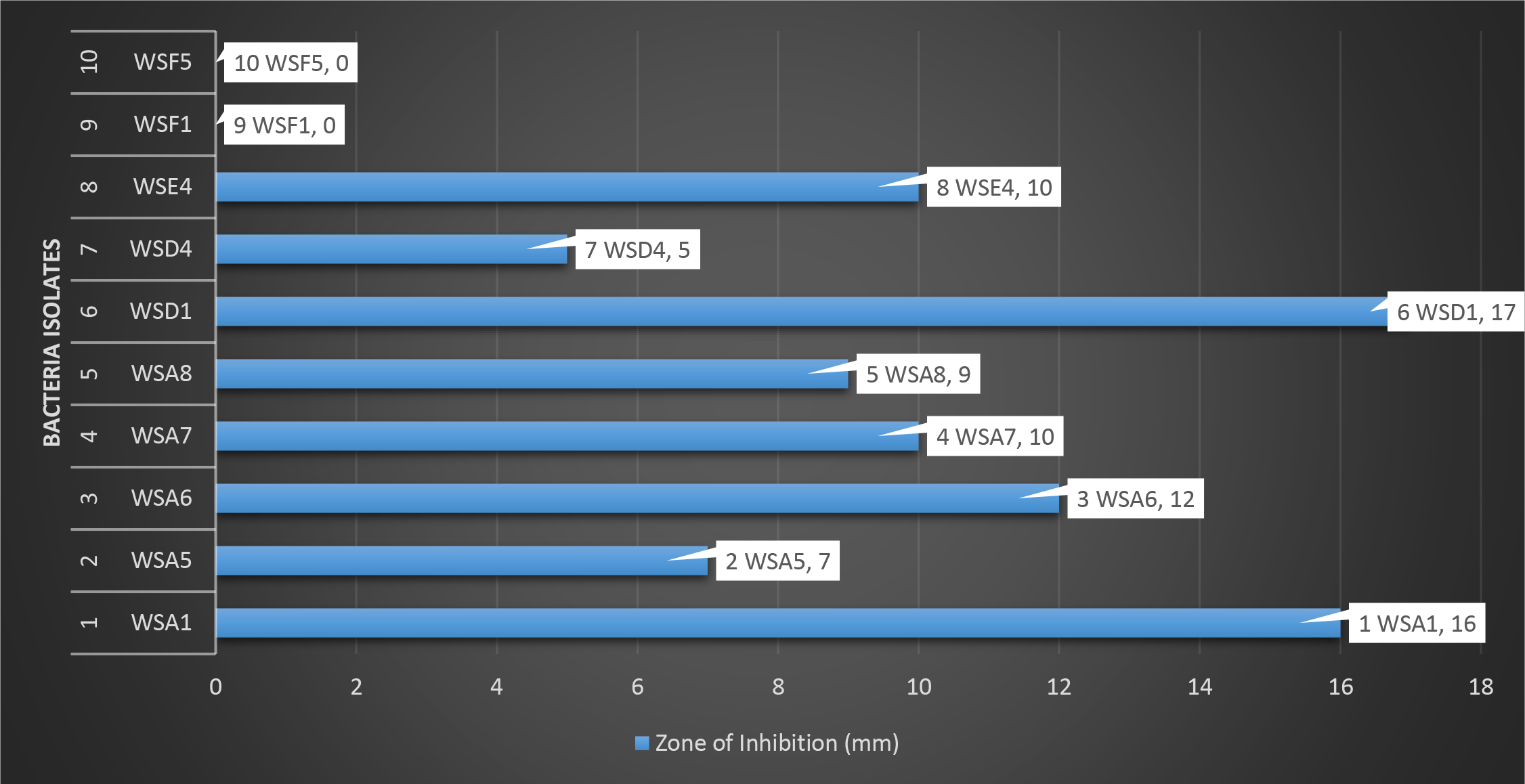
Zone of Inhibition (mm) of Isolated Bacteria on Skim Milk Plates

**Table 3:** Zone of Inhibition (cm) of Isolated Bacteria on Skim Milk Agar Plates

| S/N | Isolate Code | Zone of Inhibition (cm) | Zone of Inhibition (mm) |
| --- | --- | --- | --- |
| 1 | WSA1 | 1.6 | 16 |
| 2 | WSA5 | 0.7 | 7 |
| 3 | WSA6 | 1.2 | 12 |
| 4 | WSA7 | 1.0 | 10 |
| 5 | WSA8 | 0.9 | 9 |
| 6 | WSD1 | 1.7 | 17 |
| 7 | WSD4 | 0.5 | 5 |
| 8 | WSE4 | 1.0 | 10 |
| 9 | WSF1 | - | - |
| 10 | WSF5 | - | - |

**KEY:** - (Negative)

### 3.4 Biochemical Characterization of Bacteria Isolates

Biochemical characteristics of the isolated bacteria were presented in Table 5. The isolated bacteria exhibited different reactions to various biochemical test reagents, sugars, and physiological characteristics. They were tested for gram reaction, catalase test, indole, citrate, 6.5% NaCl, oxidase test, pH of 3.9 and 9.4, and fermentation of different sugars

**Table 4:** Screening for Proteolytic Isolates

| S/N | Isolates | Growth on Skim milk agar |
| --- | --- | --- |
| 1 | WSA1 | + |
| 2 | WSA5 | + |
| 3 | WSA6 | + |
| 4 | WSA7 | + |
| 5 | WSA8 | + |
| 6 | WSD1 | + |
| 7 | WSD4 | + |
| 8 | WSE4 | + |
| 9 | WSF1 | – |
| 10 | WSF5 | – |

**Key:** + is positive, – is negative

**Table 5:** Biochemical characterization of the Bacteria Isolated from the Waste Water

| S/N | Isolate Code | Grams | Cell Morphology | Catalase | Methyl Red | Voges Proskauer | Oxidase | Indole | Citrate | 6.5% NaCl | 20°C | 70°C | pH 3.9 | pH 9.4 | Glucose | Mannitol | Trehalose | Fructose | Raffinose | Sorbitol | Maltose | Lactose | Galactose | Bacteria Identified |
| --- | --- | --- | --- | --- | --- | --- | --- | --- | --- | --- | --- | --- | --- | --- | --- | --- | --- | --- | --- | --- | --- | --- | --- | --- |
| 1 | WSA1 | + | Rod | + | - | + | + | - | + | - | + | - | + | + | + | + | + | + | + | + | + | + | + | *Bacillus subtilis* |
| 2 | WSA5 | - | Rod | + | - | - | + | - | + | - | + | - | + | + | - | + | - | - | - | - | - | - | - | *Pseudomonas aeruginosa* |
| 3 | WSA6 | + | Rod | + | - | + | + | - | + | - | + | - | + | + | + | + | + | + | + | + | + | + | + | *Bacillus subtilis* |
| 4 | WSA7 | + | Rod | + | + | + | - | - | + | + | + | - | + | + | + | + | + | + | + | + | + | + | + | *Bacillus licheniformis* |
| 5 | WSA8 | + | Rod | + | + | + | - | - | + | + | + | - | + | + | + | + | + | + | - | + | + | + | + | *Bacillus licheniformis* |
| 6 | WSD1 | + | Rod | + | + | + | - | - | + | + | + | - | + | + | + | + | + | + | - | + | + | + | + | *Bacillus licheniformis* |
| 7 | WSD4 | - | Rod | + | - | + | + | - | + | - | + | - | + | + | - | - | - | - | - | - | - | - | - | *Pseudomonas fluorescens* |
| 8 | WSE4 | + | Rod | + | - | + | + | - | + | - | + | - | + | + | + | + | + | + | + | + | + | + | + | *Bacillus subtilis* |

**Key:** + is positive, - is negative

### 3.5 Screening for the Proteolytic Activity of the Bacteria Isolates

The results of the screening for the protease producing bacteria among the isolated bacteria were presented in Table The potential of the isolated bacteria to grow on a protease substrate was determined. Eight out of the ten isolates were able to hydrolyzed medium containing protease as its sole carbon, nitrogen, sulphur and energy source.

## 4.0 Discussion

Proteases, also known as proteolytic enzymes, play a crucial role in various biological processes and have significant industrial applications (Raveendran et al., 2018; Barrett & McDonald, 1986). These enzymes are highly valuable as they break down complex protein compounds into amino acids and peptides (Gupta et al., 2002; Verma et al., 2011). Protease-producing microorganisms have been isolated from diverse environments, including rhizosphere soil, slaughterhouse soil or floor washing, sewage, and food waste (Jadhav et al., 2020; Hakim et al., 2018; Ash et al., 2018; Prajapati et al., 2017). They have also been found in saline environments such as marine environments, sea sediments, hypersaline lakes, salted food, and soda lakes (Maruthiah et al., 2016; Boughachiche et al., 2016; Ibrahim et al., 2019). Notable bacterial producers of proteolytic enzymes include *Pseudomonas spp*., *Bacillus spp*., *Staphylococcus spp*., and *Aeromonas spp*. (Saha et al., 2011; Masi et al., 2017b). Additionally, fungal species such as *Aspergillus niger*, *Aspergillus oryzae*, *Aspergillus awamori*, *Aspergillus fumigatus*, and *Aspergillus saitoi* are known to produce extracellular acid proteases (Sielecki et al., 1991; Yongquan, 2001; Ottesen & Rickert, 1970; Shinmyo et al., 1972; Sodek & Hofmann, 1970).

Wastewater serves as a rich source of diverse microorganisms, including protease-producing bacteria (Raj et al., 2012; Sivaprakasam et al., 2011). These bacteria have adapted to thrive in the complex and nutrient-rich environment of wastewater, encountering various organic compounds, including proteins. Therefore, screening and isolating protease-producing bacteria from wastewater samples present an opportunity to discover novel microbial strains with robust proteolytic capabilities.

In this study, ten bacteria isolates were isolated from wastewater and cultured on a general-purpose medium, specifically nutrient agar. Among these isolates, eight strains exhibited growth on a medium where protease substrate served as the sole source of carbon and nitrogen. After an incubation period of 24 hours, the isolates were subjected to screening for protease activity using skim milk agar plates. The level of protease production by a microorganism can be correlated with the observed zone of hydrolysis on skim milk agar, as mentioned by Vermelho et al. (1996). This relationship indicates that a larger zone of hydrolysis on the agar plate corresponds to a higher intensity of protease production by the microorganism. This method provides a qualitative measure of protease activity and is commonly used in research and industrial settings to assess the proteolytic potential of microorganisms.

Proteolytic bacteria were identified using a combination of cell morphology, colony morphology, and biochemical tests. Specifically, *Bacillus subtilis* was identified in isolates WSA1, WSA6, and WSE4, while *Bacillus licheniformis* was determined in isolates WSA8, WSD1, and WSA7. *Pseudomonas aeruginosa* was identified in isolate WSA5, and *Pseudomonas fluorescens* in isolate WSD4. The predominance of *Bacillus spp.* among the isolates aligns with previous studies that have highlighted the efficacy of *Bacillus spp.* in producing extracellular proteases (Alnahdi, 2012; Verma et al., 2011). Table 6 shows the identification of proteolytic bacteria isolates based on combined cell morphology, colony morphology, and biochemical tests.

**Table 6:** Identification of Proteolytic Bacteria Isolates Based on Combined Cell Morphology, Colony Morphology, and Biochemical Tests

| S/N | Isolate Codes | Bacteria Identified |
| --- | --- | --- |
| 1 | WSA1 | *Bacillus subtilis* |
| 2 | WSA5 | *Pseudomonas aeruginosa* |
| 3 | WSA6 | *Bacillus subtilis* |
| 4 | WSA7 | *Bacillus licheniformis* |
| 5 | WSA8 | *Bacillus licheniformis* |
| 6 | WSD1 | *Bacillus licheniformis* |
| 7 | WSD4 | *Pseudomonas fluorescens* |
| 8 | WSE4 | *Bacillus subtilis* |

After screening ten bacterial isolates, the clearance zones were measured to assess the protease activity of each isolate. The clearance zone represents the area where the proteolytic activity of the bacteria breaks down the surrounding substrate. The clearance zone size is indicative of the effectiveness and potency of the protease produced by the bacteria. Table 3 shows the zones of inhibition (cm and mm) of isolated bacteria on skim milk agar media. Among the isolates, the highest clearance zone was observed with WSD1, identified as *Bacillus licheniformis*, measuring 1.7 cm (17mm). This indicates a strong proteolytic activity exhibited by this particular isolate. Following closely behind was WSA1, identified as *Bacillus subtilis*, with a clearance zone measuring 1.6 cm (16 mm). This suggests a substantial protease production capability. Next in line was WSA6, also identified as *Bacillus subtilis*, with a clearance zone measuring 1.2 cm (12 mm). Although slightly smaller than the previous two isolates, it still displayed a noteworthy proteolytic activity. WSE4, another *Bacillus subtilis* isolate, demonstrated a clearance zone of 1.0 cm (10 mm), showcasing its moderate protease production capacity. Both WSA7 and WSA8, identified as *Bacillus licheniformis*, exhibited clearance zones measuring 1.0 cm (10 mm) and 0.9 cm (9 mm), respectively. These isolates displayed similar protease activity, suggesting comparable proteolytic potential. Moving further down the list, WSA5, identified as *Pseudomonas aeruginosa*, produced a clearance zone of 0.7 cm (7mm). Although smaller than the previous isolates, it still demonstrated some proteolytic activity. On the other hand, WSD4, identified as *Pseudomonas fluorescens*, displayed a smaller clearance zone of 0.5 cm (5 mm), indicating a relatively lower protease production capability. Lastly, both WSF1 and WSF5 showed negative results, implying no observable protease activity in these isolates. The highest clearance zones were observed with WSD1 (1.7 cm = 17mm) identified as *Bacillus licheniformis*, WSA1 (1.6 cm = 16 mm) identified as *Bacillus subtilis*, and WSA6 (1.2 cm = 12 mm) also identified as *Bacillus subtilis* indicating their strong protease production. These results align with the study by Sidra et al. (2006), where the bacterium that exhibited the largest zone of inhibition was selected as the best producer. These findings are consistent with the research by Haider & Sanaa (2014), who isolated *Bacillus spp.* from soil and identified a strain of *Bacillus licheniformis* as the highest protease producer. Therefore, based on the results of this particular study, *Bacillus licheniformis* can be considered the best protease-producing bacterium among the tested *Bacillus spp*. However, it is important to note that the selection of the best protease-producing bacterium can depend on various factors, including the specific application and desired characteristics of the protease. Further studies and evaluations are necessary to determine the best protease producer in different contexts.

In this particular study*, Bacillus spp.* were isolated from wastewater, which aligns with previous research indicating the successful use of wastewater sludge for the production of alkaline protease using *Bacillus licheniformis* (Bezawada et al., 2010). The correlation between the findings of this study and the earlier research suggests that *Bacillus licheniformis* strains present in wastewater environments and other *Bacillus spp.* possess the potential for alkaline protease production. This highlights the viability of utilizing wastewater as a source for isolating bacteria with protease production capabilities, specifically *Bacillus spp*, which have demonstrated success in alkaline protease production in related studies (Haile & Babiye, 2018; Haider & Sanaa, 2014; Chenel et al., 2008; Tyagi et al., 2002).

The presence of *Pseudomonas spp.* isolated from wastewater in this study is consistent with findings from previous research conducted by Siddharthan et al. (2016), Masi et al. (2014), Raj et al. (2012), and Sivaprakasam et al. (2011). In the present study, two *Pseudomonas spp.* were isolated: WSA5, identified as *Pseudomonas aeruginosa*, and WSD4, identified as *Pseudomonas fluorescens*. However, these strains exhibited a comparatively lower protease production capability when compared to the *Bacillus spp*. Nonetheless, a comparative study conducted by Masi et al. (2014) indicated that a specifically isolated strain of *Pseudomonas aeruginosa* demonstrated the highest protease activity after 48 and 72 hours of incubation, surpassing the protease activity of the control strain *Bacillus subtilis*. Additionally, the protein content was also observed to be highest for *Pseudomonas aeruginosa* after 48 and 72 hours of incubation. These findings emphasize that while the *Pseudomonas spp.* isolated in this study demonstrated relatively lower protease production compared to *Bacillus spp.*, specific strains within the *Pseudomonas* genus, such as *Pseudomonas aeruginosa*, have shown the potential for significant protease activity and protein content Masi et al. (2014). Therefore, further investigation and characterization of individual *Pseudomonas* strains may uncover strains with enhanced protease production capabilities.

The comparison of protease production capabilities between *Pseudomonas spp.* and *Bacillus spp.* reveals intriguing similarities and differences. While both genera exhibit a diverse range of proteases, they differ in terms of pH range, temperature tolerance, substrate specificity, and industrial applications (Solanki et al., 2021; Razzaq et al., 2019; Zhu et al., 2013. Understanding these characteristics is essential for harnessing the potential of these microorganisms and their enzymes in biotechnological and industrial applications. Further research on specific strains within these genera will provide deeper insights into their protease production capabilities, facilitating the development of tailored enzymatic solutions for various industrial needs.

### 4.1 Limitation of the Study

The main limitation of this study is that it focuses solely on the isolation and identification of protease-producing bacteria from wastewater samples. While this provides valuable insights into the presence and potential protease production capabilities of the isolated bacteria, it does not explore other aspects of their enzymatic properties or their application in specific industrial processes. Further investigations beyond isolation and identification are necessary to fully understand the protease production capabilities of these bacteria. This may involve characterizing the specific protease enzymes produced, optimizing their production conditions, and evaluating their enzymatic properties such as pH and temperature optima, substrate specificity, stability, and potential industrial applications. Additionally, the study is limited to a specific set of samples and does not account for potential variations in protease-producing bacteria across different wastewater sources or geographic locations.

## 5.0 Conclusion

This study focused on the isolation and characterization of protease-producing bacteria from wastewater samples. The findings highlighted the presence of diverse microorganisms with proteolytic capabilities in wastewater environments. *Bacillus spp.*, particularly *Bacillus licheniformis* and *Bacillus subtilis*, demonstrated strong protease production capabilities, as evidenced by their significant clearance zones on skim milk agar plates. These results align with previous studies emphasizing the effectiveness of *Bacillus spp.* in extracellular protease production. While *Pseudomonas spp.*, including *Pseudomonas aeruginosa* and *Pseudomonas fluorescens*, exhibited relatively lower protease production compared to *Bacillus spp.*, specific strains within the *Pseudomonas* genus, have demonstrated notable protease activity and protein content in previous comparative studies. Further exploration and characterization of individual *Pseudomonas* strains may uncover potential candidates with enhanced protease production capabilities. The study also highlighted the significance of wastewater as a source for isolating bacteria with protease production capabilities. The correlation with previous research further supported the utilization of wastewater and wastewater sludge as viable resources for isolating and identifying protease-producing microorganisms, particularly *Bacillus* spp.

Overall, this study contributes to the understanding of protease-producing bacteria in wastewater environments and highlights the potential of *Bacillus spp*, especially *Bacillus licheniformis*, as strong protease producers. Further research on specific strains within *Bacillus* and *Pseudomonas* genera, as well as other microbial sources, will provide valuable insights for developing enzymatic solutions tailored to industrial applications that require proteolytic activities.

## 6.0 Recommendation

Based on the results and discussion presented in this study, it is recommended to further explore the potential of *Bacillus licheniformis* as a robust protease-producing bacterium. *Bacillus licheniformis* strains, isolated from wastewater in this study, exhibited a strong protease production capability, as evidenced by the largest clearance zone observed (1.7 cm = 17mm) compared to other tested strains. This finding aligns with previous research that highlights the effectiveness of *Bacillus licheniformis* in extracellular protease production (Sidra et al., 2006; Haider & Sanaa, 2014).

Further investigations should focus on characterizing the proteases produced by *Bacillus licheniformis* strains, including their biochemical properties, substrate specificity, and potential industrial applications. Additionally, optimization studies, such as medium composition and culture conditions, can be conducted to enhance protease production yields. The isolation of *Bacillus licheniformis* from wastewater samples suggests the possibility of utilizing this bacterium in wastewater treatment processes, where protease activity can aid in the degradation of organic compounds.

It is also recommended to explore specific strains within the *Pseudomonas* genus, particularly *Pseudomonas aeruginosa*, for their protease production capabilities. Although the *Pseudomonas spp.* isolated in this study demonstrated relatively lower protease production compared to *Bacillus spp*, previous research has indicated the potential of certain *Pseudomonas* strain, to exhibit significant protease activity (Masi et al., 2014). Further investigation and characterization of individual *Pseudomonas* strains may unveil strains with enhanced protease production capabilities, enabling their utilization in specific industrial applications.

To fully exploit the protease production potential of both *Bacillus spp.* and *Pseudomonas spp*., a comparative study can be conducted to evaluate their enzymatic properties, including pH and temperature optima, stability, substrate specificity, and compatibility with different industrial processes. This would provide valuable insights into the selection and application of the most suitable protease-producing microorganism for specific industrial sectors such as detergent manufacturing, food processing, pharmaceuticals, and waste management.

In conclusion, the findings of this study underscore the significance of *Bacillus licheniformis* and other *Bacillus spp.* as a promising protease producer, particularly when isolated from wastewater sources. The potential of *Pseudomonas spp*, specifically *Pseudomonas aeruginosa*, should also be explored further. Understanding the enzymatic capabilities of these microorganisms and their proteases will pave the way for the development of tailored enzymatic solutions and biotechnological applications in various industries.

## Acknowledgements

The authors would like to express their sincere gratitude to Dr. K.O Awojobi and Dr. S.M. Adeyemo for their invaluable supervision and support throughout the research project. The authors are also grateful to the laboratory assistants for their unwavering technical support, which greatly contributed to the successful completion of the project. Special thanks go to all the dedicated staff members of the Department of Microbiology at Obafemi Awolowo University, Nigeria, for their invaluable assistance and collaboration. Their contributions played a crucial role in the overall accomplishment of this study.

## Contribution of authors

AFQ and LRA both contributed to various aspects of the study. AFQ was responsible for material preparation, data collection, microbiological analysis, data analysis and interpretation, literature searches, and writing the initial draft of the manuscript. LRA, on the other hand, participated in the sampling process and conducted microbiological analysis of the samples. Both authors were involved in reviewing the manuscript draft, and all authors read and approved the final version of the manuscript.

## Data Availability

The corresponding author can provide the data used to support the findings of this study upon request.

## Source of funding

The study did not receive any funding.

## Conflict of interest

The authors affirm that they have no conflicts of interest to declare.

## Notes

### Competing Interest Statement

The authors have declared no competing interest.

